# Liver sinusoidal endothelial cells integrate metabolic and immune signals for MAPK-dependent BMP6 regulation and hepcidin induction

**DOI:** 10.64898/2026.05.07.723498

**Authors:** Ruiyue Qiu, Stefania Cucinelli, Christina Mertens, Silvia Colucci, Sandro Altamura, Matthias W. Hentze, Martina U. Muckenthaler

## Abstract

Liver sinusoidal endothelial cells (LSECs) separate the blood from the hepatic parenchyma and thus are at the frontline as scavengers of blood-borne waste, pathogens and metabolic stimuli. LSECs are also critical for sensing systemic iron availability by controlling the synthesis of bone morphogenetic protein (BMP) 6, which is essential for hepcidin expression in hepatocytes. Hepcidin maintains systemic iron homeostasis by inhibiting dietary iron uptake and iron release from iron recycling macrophages. Hepcidin is also an acute-phase protein and its activation by inflammation requires active BMP signaling. It is incompletely understood how signals derived from inflammation, cellular damage and iron are integrated by the liver to assure adequate hepcidin expression. Here, we show that *Bmp6* expression is activated in primary LSEC cultures upon their exposure to damage-associated molecular patterns (DAMPs), such as heme and myoglobin, pathogen-associated molecular patterns (PAMPs), such as lipopolysaccharide (LPS) and Fibroblast-Stimulating Lipopeptide-1 (FSL1), or oxidative stress inducers (H_2_O_2_). All regulatory cues converge at the MAPK signaling pathway, although the specific signaling branches involved are stimulus-specific. Notably, *Bmp6* upregulation in LSECs in response to all signals tested is strongly enhanced by the hepatocyte secretome. As hepatocytes critically depend on active BMP/SMAD signaling to control hepcidin activation, our results reveal that multiple sources of signaling input activating *Bmp6* in LSECs and hepcidin in hepatocytes serve to determine BMP/SMAD signaling strength. Furthermore, our findings identify hypoferremia, the result of high hepcidin levels due to elevated *Bmp6*, as a convergent response in conditions of inflammation, oxidative stress and cellular damage.

**Highlights:**

- DAMPs (heme and myoglobin), PAMPs (LPS) and oxidative stress activate *Bmp6* mRNA expression via the MAPK signaling pathway
- The TLR/MAPK/BMP6 regulatory axis integrates inflammatory and iron signals
- Our work uncovers a novel connection between innate immune sensing, oxidative stress and hepatic iron homeostasis

## Introduction

Control of systemic iron homeostasis is a dynamic process that ensures adequate iron availability for cellular processes, such as oxygen transport, mitochondrial respiration or DNA synthesis, while preventing the formation of reactive oxygen species (ROS), and consequent oxidative damage and ferroptosis (1). In mammals, this balance is achieved at the systemic level by the hepcidin–ferroportin (FPN, SLC40A1) regulatory axis, that maintains iron levels within physiological limits. Hepcidin is synthesized and secreted by hepatocytes and inhibits iron fluxes via FPN from duodenal enterocytes, macrophages, and hepatocytes into the blood (1). The functioning of this hormonal axis critically depends on cellular regulatory systems that control the expression levels of hepcidin. A fundamental and incompletely understood question is how the liver integrates signals derived from inflammation, oxidative stress and cell damage with iron availability to control hepcidin expression.

Hepcidin transcriptional activation critically depends on the bone morphogenetic protein (BMP)-SMAD signaling pathway. BMPs are multifunctional members of the transforming growth factor-β (TGF-β) superfamily that activate hepcidin expression *in cell lines* or *ex vivo* (2-5). *In vivo*, BMP2 and BMP6 are the major regulators of the *Hepcidin* gene expression (6-8). BMP2 and BMP6 are produced predominantly by liver sinusoidal endothelial cells (LSECs) where they act in a paracrine manner on hepatocytes. There, they bind to BMP type I (ALK2/ALK3) and type II (ACVR2A/BMPR2) receptors to activate the SMAD1/5/8 pathway that controls *Hepcidin* transcription (1).

In addition to dietary iron uptake and recycling of senescent erythrocytes by macrophages, hemolysis represents an important physiological and pathological source of circulating iron. During red blood cells (RBCs) breakdown, large amounts of heme and iron are released into the bloodstream. Under conditions of limited hemolysis, free hemoglobin and heme are rapidly scavenged by plasma proteins such as haptoglobin and hemopexin, respectively (9). However, during excessive hemolysis, caused by infection, inflammation, or in the context of genetic disorders such as sickle cell disease (SCD), these buffering systems are overwhelmed, leading to increased levels of unbound heme and iron (9, 10). Free heme acts as a potent damage-associated molecular pattern (DAMP), triggering toll-like receptor 4 (TLR4) signaling and oxidative stress responses (11, 12).

LSECs are specialized endothelial cells separating the blood stream and hepatic parenchyma, and are therefore among the first cells in the liver to encounter inflammatory cues, DAMPs and PAMPs (13). This setting makes them not only crucial sensors of systemic iron levels but also key responders to hemolytic and inflammatory stress. LSECs actively participate in the clearance of spleen-derived hemolytic products, such as hemoglobin and erythrocytic membrane (14). Moreover, in SCD LSECs scavenge hemoglobin, and their impairment by intracellular accumulation of sickling hemoglobin aggravates liver injury in SCD mice (15). Moreover, in the context of liver disease, such as non-alcoholic steatohepatitis (NASH), LSECs acquire a proinflammatory phenotype and release cytokines and chemokines to recruit immune cells, thus exacerbating inflammation (13). Understanding how LSECs integrate these signals to control *Bmp6* mRNA expression is essential for linking iron metabolism to immune and stress pathways in the liver.

How *Bmp2* and *Bmp6* mRNA expression in LSECs is controlled by iron remains under discussion. Some studies proposed that *Bmp6* upregulation in LSECs requires intracellular iron accumulation, e.g. from uptake of non-transferrin-bound iron (NTBI). NTBI accumulation and the resulting oxidative stress activate nuclear factor erythroid 2-related factor 2 (NRF2) (16, 17) and c-JUN (18), which then induce *Bmp6* transcription in a cell-autonomous manner. Alternatively, iron-mediated *Bmp6* induction in primary cultured LSECs was shown to occur in an NRF2-independent manner via the transcription factor ETS1, which signals downstream of the p38/JNK mitogen-activated protein kinase (MAPK) pathway (19). Moreover, it has been recently demonstrated that target of rapamycin complex 2 (mTORC2) acts as an iron sensor in LSECs. Mechanistically, iron mediates mTORC2 degradation, leading to forkhead box protein O1 (FOXO1)-dependent *Bmp6* upregulation (20).

The model that intracellular iron accumulation by LSECs triggers increased *Bmp6* mRNA levels was challenged by findings in mouse models of hemochromatosis (HC). For example, LSECs from *Hjv*-KO and Fpn(C326S) mice show elevated *Bmp6* mRNA expression despite a cellular molecular signature of iron deficiency (21, 22). Moreover, *Bmp6* mRNA levels increase during aging, despite an unchanged iron content in non-parenchymal cells, while iron deposition is rather localized in hepatocytes (19). Importantly, iron uptake into LSECs via transferrin receptor 1 (TFR1) or the NTBI transporter ZIP8 does not appear to be essential for controlling BMP2 and BMP6 *in vivo*, as the conditional deletion of either in LSECs does not significantly affect systemic iron homeostasis (16, 23, 24). Similarly, the LSEC-specific deletion of the abundantly expressed HC protein HFE leaves iron homeostasis unaffected (25). By contrast, LSEC-specific *Fpn*-KO mice increase *Bmp6* mRNA levels (26). Thus, the iron-sensing process may involve additional liver cell types.

Our previous work provided evidence that the iron-dependent *Bmp6* control in LSECs is driven by a paracrine signaling mechanism, in which hepatocyte-derived factors enhance iron-dependent *Bmp6* induction (22). However, the identity of the signaling pathways and mediators involved in this cross-talk remained unclear.

Here we identify the MAPK signaling pathway as a regulatory hub that controls *Bmp6* mRNA expression not only to iron-related signals but also to oxidative stress (H_2_O_2_) and inflammatory stimuli (heme or LPS). With the exception to iron, these signals trigger TLR4 activation and induce different branches of the MAPK pathway. Collectively, our results highlight both shared and stimulus-specific pathways involved in *Bmp6* control and reveal a novel connection between innate immune sensing, oxidative stress and hepatic iron homeostasis. Additionally, we show that hepatocyte-secreted factors modulate the responsiveness of LSECs to all external signals tested, further highlighting the importance of intrahepatic cell-to-cell communication in iron regulation.

## Methods

### Mice

C57BL/6N wild-type mice were housed under specific pathogen-free (SPF) conditions at the Interfakultäre Biomedizinische Forschungseinrichtung (IBF) animal facility, Heidelberg University (Germany). All mice were kept on a standard 12-hour light/dark cycle and fed a standard rodent diet containing 200 ppm iron, with ad libitum access to food (LASQCdiet® Rod18-A, LASvendi, Soest, Germany) and water. All animal breeding and experimental procedures were conducted following institutional and governmental regulations and were approved by the Regierungspräsidium Karlsruhe (T-47/22, T-52/23, T-48/24).

### Isolation and culture of primary LSECs and hepatocytes

Primary mouse LSECs and hepatocytes were isolated as previously described with minor modifications (27). Details about primary cell isolation and cell culture treatments are described in the supplementary methods section.

### Protein extraction and Western blotting analysis

Protein extraction was performed as described in the supplementary section. Equal amounts of protein were separated by 10% SDS-PAGE and subsequently transferred for Western blotting using the antibodies listed in Table S1. Chemiluminescence signals were captured and quantified using the Vilber Lourmat Fusion-FX imaging system.

### Quantitative Real-Time PCR (RT-qPCR)

Total RNA was extracted from cells using RNA-Mini Kits (Bio&Sell # BS67.311.0250) according to the manufacturer’s instructions. RNA was reverse-transcribed using random hexamer primers and the RevertAid H Minus Reverse Transcriptase (Life Technologies #EP0452). RT-qPCR was performed using SYBR Green on a StepOnePlus Real-Time PCR System (Applied Biosystems) with the primers listed in Table S2. Results were calculated using the Pfaffl method (28).

### RNA sequencing (RNA-seq)

Strand-specific RNA-seq libraries were prepared using the The NEBNext UltraExpress RNA Library Prep Kit and sequenced in single-end mode on a platform of Element AVITI (150 cycles Kit). Details of the RNA-seq data analysis are reported in the supplementary information. RNA-seqdata are available in Biostudies (Arrayexpress) under the accession number E-MTAB-1637.

### Statistics analysis

Data are presented as mean ± standard deviation (SD). Statistical analyses were performed using GraphPad Prism version 8. Two-tailed Student’s t-tests, one-way ANOVA, or two-way ANOVA were used where appropriate. Statistical significance was defined as p < 0.05 (*), p < 0.01 (**), p < 0.001 (***), and p < 0.0001 (****).

## Results

### The TLR4/MAPK regulatory axis controls *Bmp6* mRNA expression in primary LSECs in presence of hepatocyte secretome

TLRs orchestrate immune responses and cellular activation by detecting PAMPs and DAMPs. LSECs express multiple TLRs, enabling them to sense microbial components and endogenous damage signals. Combined with their efficient scavenger function, this makes LSECs sensitive sentinel cells that initiate acute-phase responses in the liver (13). To investigate whether the activation of TLRs in LSECs can control iron homeostasis via *Bmp6*, we first tested LPS, a membrane component of gram-negative bacteria, as trigger of the innate immune response with reported functions as TLR4 ligand in LSECs (29). Based on our previous findings showing that *Bmp6* mRNA upregulation by iron is enhanced in presence of hepatocyte conditioned medium (CM) in primary cultured LSECs (22), we evaluated the effect of LPS in presence or absence of CM. In addition, we tested the TLR4 ligand heme, a DAMP released during hemolytic processes, such as SCD or sepsis (11, 12). Both LPS and heme required CM to activate *Bmp6* mRNA expression (Figure 1A).

**Figure 1.**
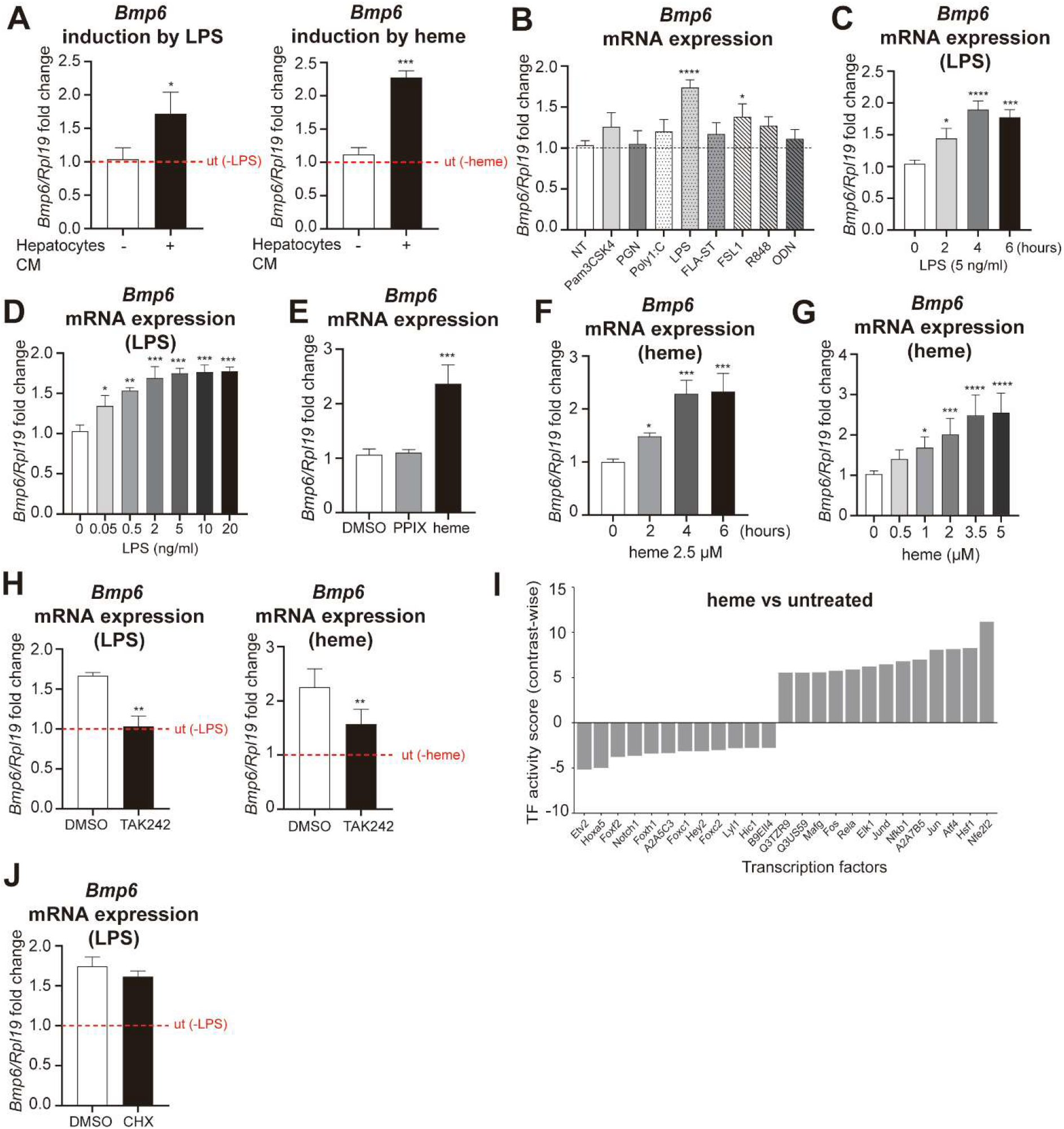
LPS and heme induce *Bmp6* expression in primary LSECs via TLR4. (A) *Bmp6* mRNA expression in primary LSECs treated with 5 ng/mL LPS or 2.5 μM heme for 6 h in the presence or absence of hepatocyte-conditioned medium. (B) *Bmp6* mRNA expression in primary LSECs treated with various TLR ligands: Pam3CSK4 (TLR1/2), PGN (TLR2), Poly I:C (TLR3), LPS (TLR4), FLA-ST (TLR5), FSL1 (TLR2/6), R848 (TLR7/8), and ODN (TLR9) or vehicle control (NT, non-treated) for 6 h. (C-D) *Bmp6* mRNA expression in primary LSECs treated with increasing concentrations of LPS for 6 h or with 5 ng/mL LPS for 2, 4, and 6 h. (E) *Bmp6* mRNA expression in LSECs treated with 2.5 μM heme or protoporphyrin IX (PPIX) for 6 h. (F-G) *Bmp6* mRNA expression in primary LSECs treated with increasing concentrations of heme for 6 h or treated with 2.5 μM heme for 2, 4, or 6 h. (H) *Bmp6* mRNA expression in primary LSECs pre-treated with TAK242 (5 μM) or DMSO for 1 h, followed by 5 ng/mL LPS or 2.5 μM heme treatment for 6 h. (I) Transcription factor activity analysis by RNA-seq of LSECs treated with heme with respect to untreated controls (contrast-wise), in presence of hepatocyte-conditioned medium. (J) *Bmp6* mRNA expression in primary LSECs pre-treated with CHX (5 μM) or DMSO for 1 h, followed by 5 ng/mL LPS treatment for 6 h. Cell culture experiments, except those in panel A, were always conducted in the presence of hepatocyte-conditioned medium. Gene expression levels were assessed by RT-qPCR, normalized to the housekeeping gene *Rpl19*, and expressed as fold change relative to vehicle-treated controls. The dashed line (ut) represents the mRNA expression of LSECs treated with the conditions shown, in the absence of LPS or heme. Data are obtained from three or four independent experiments and displayed as mean ± SD. Statistical significance: *p<0.05, **p<0.01, ***p<0.001, ****p<0.0001, one-way ANOVA or Student’s t-test. CM, conditioned medium; PPIX, protoporphyrin IX; CHX, cycloheximide.

To assess whether the activation of other TLRs induces *Bmp6* mRNA expression in presence of CM, we treated LSECs with different ligands activating TLR1 to TLR9. Among the tested ligands, only the treatment with the TLR4 agonist lipopolysaccharide (LPS) strongly increased *Bmp6* mRNA levels, while fibroblast-stimulating lipopeptide-1 (FSL1), which binds TLR2/6, induced a more modest response (Figure 1B). Of note, TLR4 is the most abundantly expressed TLR in primary cultured LSECs (Figure S1A).

We next characterized the *Bmp6* mRNA response to TLR4 activation in more detail. We show that LPS induced *Bmp6* mRNA expression as early as 2 hours after treatment, even at the very low concentration of 0.05 ng/mL (Figure 1C, 1D). Likewise, heme, but not protoporphyrin IX (PPIX), induced *Bmp6* mRNA expression in a time- and dose-dependent manner, indicating that the iron-containing core of heme is essential for this response (Figure 1E, 1F and 1G). Similarly, myoglobin, a damage signal detectable in the plasma of patients with heart diseases (30), also increased *Bmp6* mRNA levels in presence of CM (Figure S1B), suggesting that DAMPs derived from other organs (e.g. the heart) can induce responses in LSECs to control systemic iron homeostasis. Of note, LSECs were treated with 50 µM myoglobin to model elevated extracellular myoglobin levels associated with acute myocardial infarction damage (31). As BMP6 activates hepcidin expression, this finding may help to explain the iron deficiency observed in 30% of heart failure patients (32).

To confirm the role of TLR4 in the LPS, heme and myoglobin-controlled *Bmp6* response, we pre-treated LSECs with the TLR4 inhibitor TAK242. TAK242 effectively reduced *Il6* mRNA induction in response to all treatments (Figure S1C-E) and significantly, but not fully, attenuated *Bmp6* mRNA activation by all stimuli (Figure 1H, S1F). This result further supports the importance of TLR4 in the stimulus-mediated *Bmp6* response, while it also implicates contributions by other signaling networks.

To identify signaling networks involved in *Bmp6* induction following TLR4 activation, we performed transcription factor activity inference analysis on RNA-seq data from primary LSECs treated with or without heme, in presence of hepatocyte CM. Consistent with a function as TLR ligand in LSECs, heme exposure increased the activities of key inflammation-related transcription factors (NFkB1 and RELA) (Fig. 1I). Moreover, gene set enrichment analysis (GSEA) showed enriched pathways related to inflammatory signaling (Figure S1G) upon heme treatment, which consistently led to the increase in the mRNA expression of pro-inflammatory cytokines (e.g. *Il6, Tnfa, Il1b, Il12b*) and chemokines (e.g., *Cxcl10, Ccl2, Ccl4*) (Figure S1H). Similarly, we observed an induction of proinflammatory cytokines upon LPS treatment (Fig. S1I).

To exclude that *Bmp6* upregulation in response to TLR activation is secondary to the synthesis of cytokines or other effectors, we pre-treated LSECs with cycloheximide to inhibit protein synthesis, followed by LPS addition. Our data show that cycloheximide fails to reduce LPS-mediated *Bmp6* induction (Figure 1J), indicating that *Bmp6* is a primary response gene. By contrast, cycloheximide effectively suppressed LPS-induced nitric oxide synthase 2 (*Nos2*) mRNA expression, which requires *de novo* protein synthesis (33), thereby confirming its inhibitory activity (Figure S1J).

Consistent with previous observations that hemolysis impairs endothelial function (10, 12, 15), LSEC exposure to heme reduced the activity of transcription factors maintaining LSEC identity (ETV2 and NOTCH1) (Figure 1I) (34, 35). In addition, the mRNA expression of hallmark LSEC genes, including *Stab2, Cd146* (*Mcam*), *Vegfr3* (*Flt4*), *Vegfr2* (*Kdr*), *Oit3*, and *Cd32b* (*Fcgr2b*), was markedly reduced (Figure S1K).

The most upregulated pathway upon heme treatment is the oxidative stress response (Figure S1G). Indeed, the activity of the transcription factor NRF2 (Figure 1I) and mRNA expression of its target genes, including *Hmox1* and *Nqo1*, are strongly increased (Figure S1L). By contrast, myoglobin treatment does not affect *Hmox1* mRNA levels (Figure S1M) and LPS treatment even reduced *Hmox1* and *Nqo1* expression (Figure S1N), indicating that, unlike heme, LPS does not activate (but possibly suppress) the NRF2 pathway. Because different signals activating TLR4 induce a variable response of NRF2 target genes, we conclude that NRF2 is not a common regulatory hub to induce *Bmp6* mRNA expression downstream of TLR signaling.

We next focused on transcription factors acting downstream of the MAPK pathway (JUN, JUND, ELK1 and FOS) that were strongly activated following heme treatment (Figure 1I). MAPK signaling is activated by cell surface receptors such as TLRs, receptor tyrosine kinases (RTKs) and G protein-coupled receptors (GPCRs) which induce one or several signaling branches, including ERK1/2, JNK and p38 that control key cellular processes such as proliferation, differentiation, survival, and stress responses (36). In line with the transcription factor responses in our RNA-seq data analysis, we detected increased phosphorylation of JNK, ERK1/2, and p38 in LSECs following heme stimulation (Figure 2A), and LPS treatment (Figure 2B).

**Figure 2.**
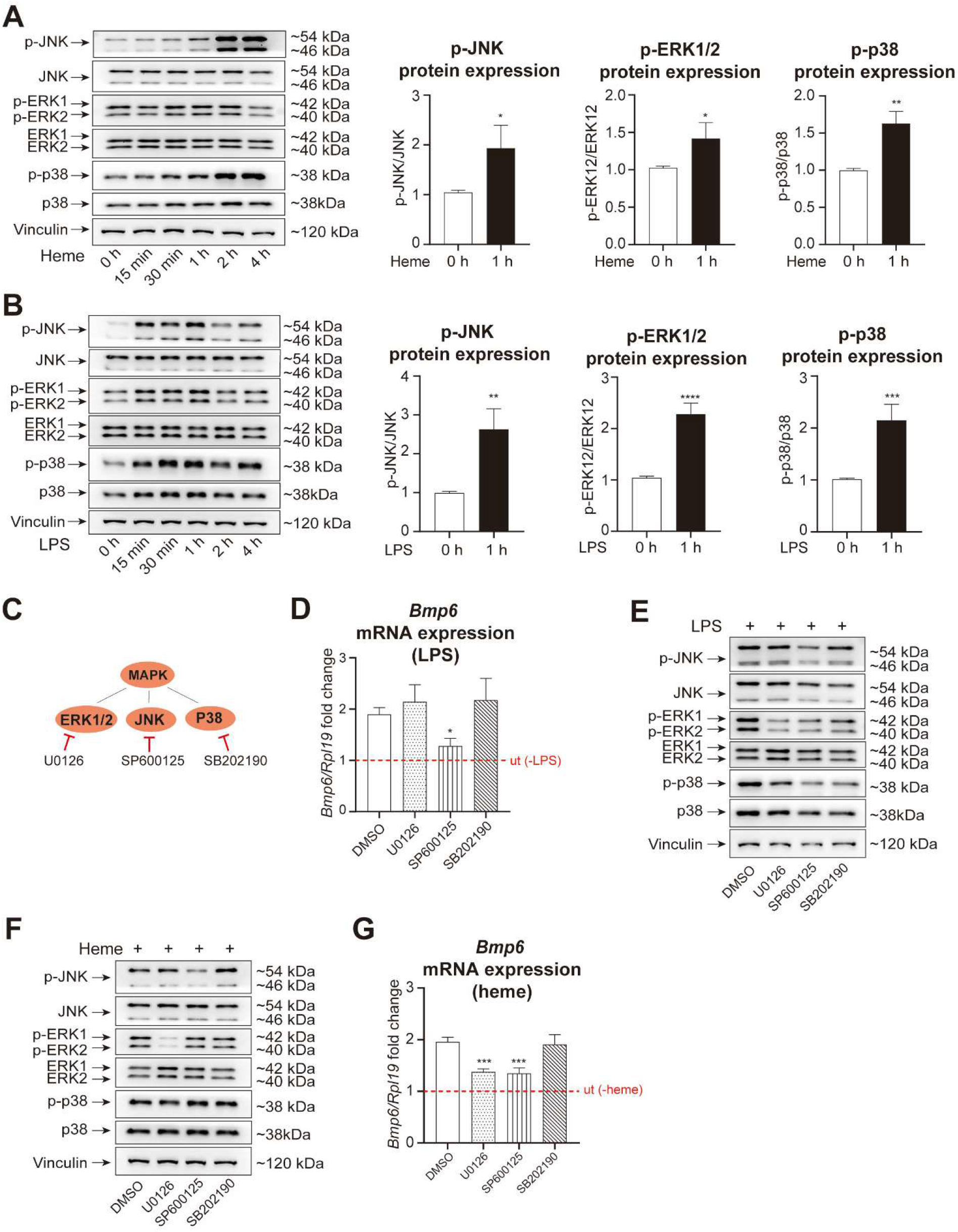
LPS and heme-induced *Bmp6* expression depend on MAPK signaling. (A-B) Western blot analysis and quantification of phosphorylated and total JNK, ERK1/2, and p38 in LSECs treated with 2.5 μM heme or 5 ng/mL LPS for the indicated time points (15 min to 4 h). (C) Scheme showing the MAPK branches inhibitors and their respective targets. (D) *Bmp6* mRNA expression in primary LSECs pre-treated with U0126 (10 μM), SP600125 (5 μM), SB202190 (10 μM), or DMSO for 1 h, followed by 5 ng/mL LPS treatment for 6 h. (E-F) Western blot analysis of phospho-JNK, JNK, phospho-ERK1/2, ERK1/2, phospho-p38, and p38 in primary LSECs pre-treated with U0126 (10 μM), SP600125 (5 μM), SB202190 (10 μM), or DMSO for 1 h, followed by 5 ng/mL LPS or 2.5 μM heme for 1 h. (G) *Bmp6* mRNA expression in primary LSECs pre-treated with the inhibitors described in panel C, followed by 5 ng/mL heme treatment for 6 h. All cell culture experiments were conducted in the presence of hepatocyte-conditioned medium. Gene expression levels were assessed by RT-qPCR, normalized to the housekeeping gene *Rpl19*, and expressed as fold change relative to vehicle-treated controls. The dashed line (ut) represents the mRNA expression of LSECs treated with the conditions shown, in the absence of LPS and heme. Data are obtained from three independent experiments and displayed as mean ± SD. Western blots in panels A-B, E-F show a representative replicate of three independent experiments. Densitometric quantification displays mean ± SD of all biological replicates. Statistical significance: *p<0.05, **p<0.01, ***p<0.001, ****p<0.0001, one-way ANOVA or Student’s t-test.

To next assess whether an activated MAPK signaling pathway controls *Bmp6* induction in LSECs, we treated primary LSECs with pharmacological inhibitors of MAPK components: U0126 (ERK1/2 via MEK1/2), SP600125 (JNK) and SB202190 (p38) (Figure 2C). Pre-treatment of primary LSECs with the JNK inhibitor SP600125 completely blocked LPS-induced *Bmp6* mRNA expression (Figure 2D). Notably, SP600125 not only inhibited JNK activation, but also suppressed LPS-induced phosphorylation of ERK1/2 and p38 (Figure 2E), suggesting off-target effects that may explain its impact on *Bmp6* mRNA expression. By contrast, U0126 and SP600125 effectively blocked heme-induced phosphorylation of ERK1/2 and JNK, respectively, while SB202190 had no significant effect (Figure 2F). Inhibition of ERK1/2 and JNK, but not p38, significantly reduced *Bmp6* and *Nqo1* mRNA expression following heme treatment (Figure 2G and Figure S1O), suggesting that MAPK signaling is critical for both *Bmp6* induction and NRF2 pathway activation.

Taken together, our results demonstrate that TLR4 activation induces *Bmp6* mRNA expression in LSECs via MAPK signaling, a mechanism shared by both PAMPs (LPS) and DAMPs (heme). Interestingly, the signaling branches involved in *Bmp6* activation are signal-specific.

### Iron-induced *Bmp6* mRNA expression requires the activation of MAPK signaling pathway in LSECs

We next tested whether the MAPK pathway is further required for activating *Bmp6* mRNA expression in primary LSEC cultures maintained with hepatocyte CM with and without iron, as previously established (22). Our data show that iron treatment of primary LSECs increased the levels of phosphorylated ERK1/2 in a time-dependent manner. This finding is consistent with the fact that FeNTA induces ROS accumulation, which can activate ERK1/2 (37). Other branches of the MAPK pathway remained unaffected (JNK) or showed decreased phosphorylation (p38) (Figure 3A). Pharmacological inhibition of the ERK1/2-MAPK pathway targeting RAF (Raf265), MEK1/2 (U0126) and ERK1/2 (Ulixertinib) reduced ERK1/2 phosphorylation (except Ulixertinib) (Figure 3B), downregulated the MAPK target gene *Ccnd1* (Figure S2A) and, importantly, reduced iron-induced *Bmp6* mRNA expression (Figure 3C). This confirms a critical role for the ERK1/2 pathway in controlling *Bmp6* mRNA levels in response to iron in presence of hepatocyte CM. Ulixertinib failed to diminish ERK1/2 phosphorylation, as it blocks the downstream activity of ERK1/2 without affecting its phosphorylation, as previously observed (38). Notably, MAPK inhibition also abolished the iron-induced transcription of the two downstream targets of NRF2 *Hmox1* and *Nqo1* (Figure S2B). These data suggest that MAPK signaling contributes to the regulation of *Bmp6* and well-known NRF2 targets in LSECs.

**Figure 3.**
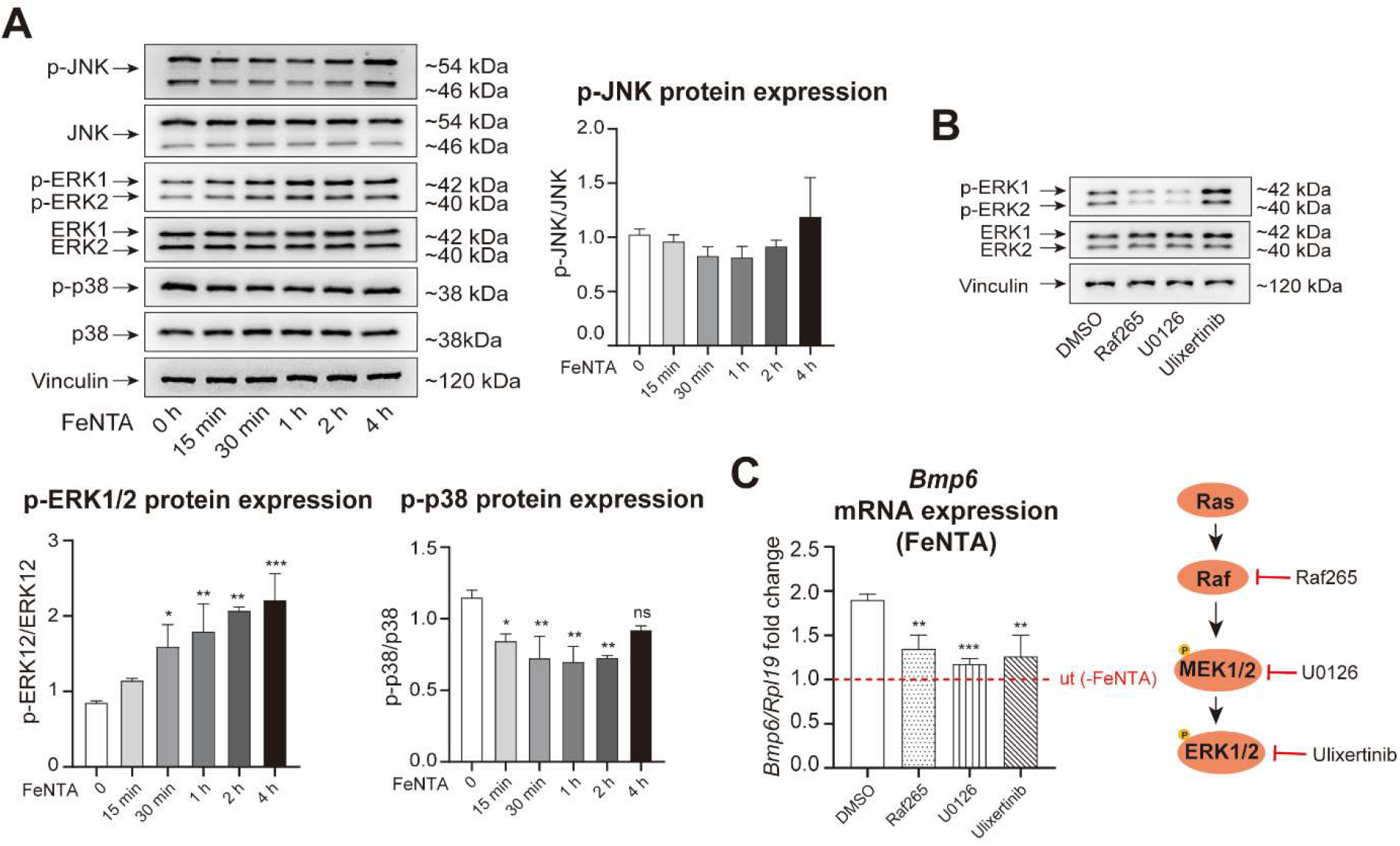
The MAPK signaling pathway is involved in iron-induced *Bmp6* expression in LSECs in the presence of hepatocyte-conditioned medium. (A) Western blot analysis and quantification of phospho-JNK, JNK, phospho-ERK1/2, ERK1/2, phospho-p38, and p38 in primary LSECs stimulated with 50 μM FeNTA. (B) Western blot analysis of phospho-ERK1/2 and total ERK1/2 in primary LSECs pre-treated with RAF265, U0126, Ulixertinib, or DMSO as in (C), followed by 50 μM FeNTA for 1 h. (C) *Bmp6* mRNA expression in primary LSECs pre-treated with RAF265 (5 μM), U0126 (10 μM), Ulixertinib (10 μM), or DMSO for 1 h, followed by 50 μM FeNTA for 6 h. All cell culture experiments were conducted in the presence of hepatocyte-conditioned medium. Data in C are RT-qPCR results, normalized to the housekeeping gene *Rpl19* and presented as fold change relative to vehicle controls. The dashed line (ut) indicates the *Bmp6* mRNA expression of LSECs treated with DMSO or the inhibitors in the absence of FeNTA. Data are derived from three independent experiments and presented as mean ± SD. Western blots in panels A-B show a representative replicate of three independent experiments. Densitometric quantification displays mean ± SD of all biological replicates. Statistical significance: *p<0.05, **p<0.01, ***p<0.001, ****p<0.0001, one-way ANOVA or Student’s t-test. FeNTA, ferric nitrilotriacetate.

### Oxidative stress induced by H_2_O_2_ controls *Bmp6* mRNA expression in LSECs via MAPK signaling in presence of hepatocyte CM

Given that iron is a potent inducer of oxidative stress, we next assessed ROS production in LSECs by iron compared to H_2_O_2_ treatment. We show that FeNTA treatment generated a stronger and longer-lasting ROS signal than H_2_O_2_-induced oxidative stress (Figure 4A). Interestingly, also H_2_O_2_ treatment increased *Bmp6* mRNA expression only in the presence of hepatocyte CM (Figure 4B). ROS accumulation and *Bmp6* mRNA induction in response to H_2_O_2_ were effectively prevented by the ROS scavenger N-Acetyl-L-Cysteine (NAC) (Figure 4A, C). H_2_O_2_ also increased the mRNA levels of the NRF2 target genes *Hmox1* and *Nqo1*, although to a lesser extent than FeNTA at 6 h, a response prevented by NAC treatment (Figure S3A). While NAC completely abolished the *Bmp6* response to H_2_O_2_, it only partially reduced iron-induced *Bmp6, Hmox1* and *Nqo1* mRNA expression (Figure 4C; S3A). Notably, NAC treatment was not able to fully restore ROS basal levels in FeNTA-treated LSECs. We therefore cannot exclude the involvement of oxidative stress-dependent mechanisms in controlling FeNTA-mediated *Bmp6* response.

**Figure 4.**
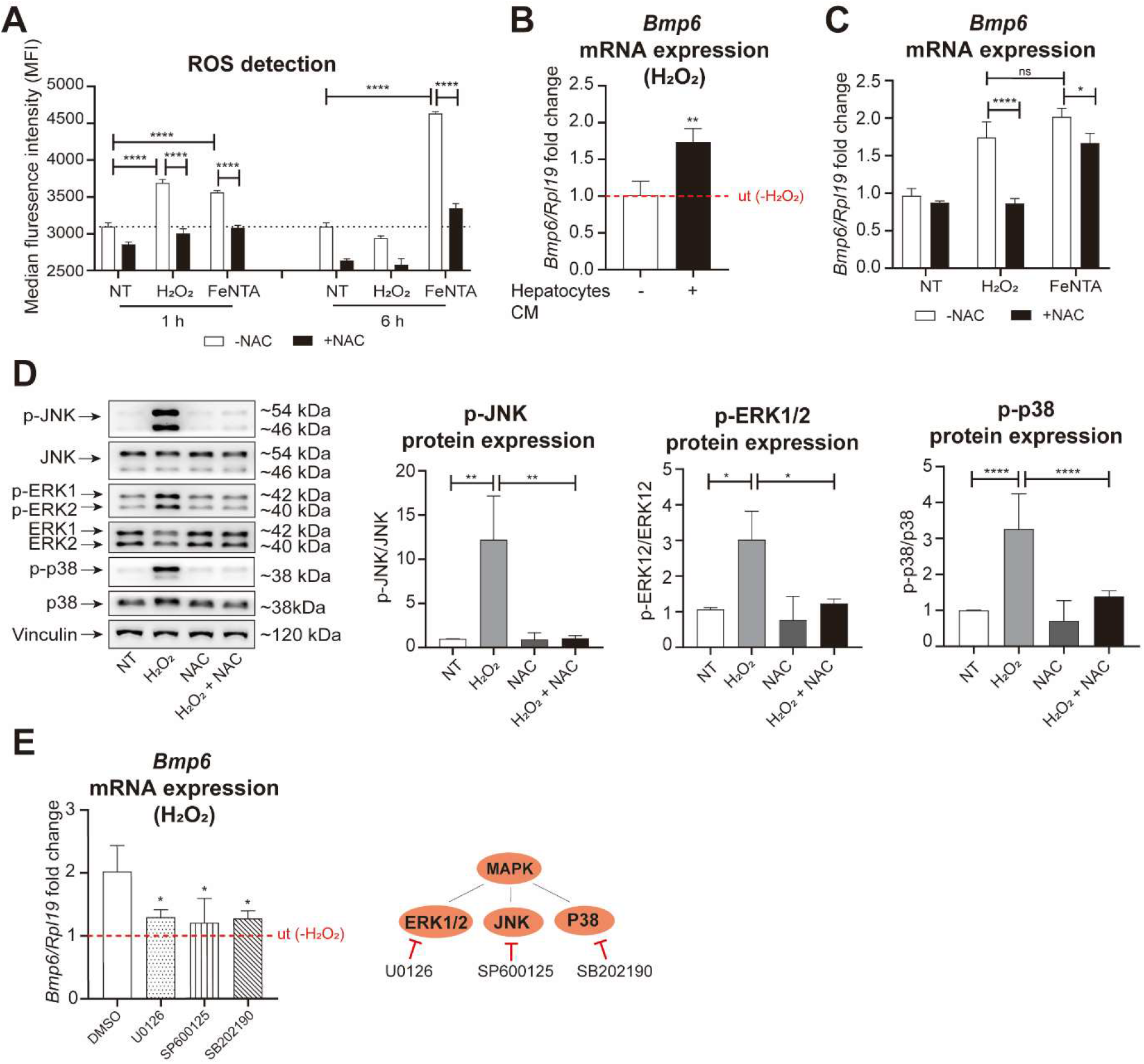
Oxidative stress-induced *Bmp6* expression depends on MAPK signaling pathway in the presence of hepatocyte-conditioned medium. (A) Measurement of ROS content in primary LSECs treated with H_2_O_2_ (100 μM) or FeNTA (50 μM) for 1 or 6 h in the absence or presence of NAC (5 mM), analyzed by flow cytometry. (B) *Bmp6* mRNA expression in primary LSECs treated with 100 μM H_2_O_2_ or vehicle control (NT, non-treated) for 6 h in the presence or absence of hepatocyte-conditioned medium. (C) *Bmp6* mRNA expression in primary LSECs pre-treated with NAC (5 mM) for 1 h, followed by H_2_O_2_ (100 μM) or FeNTA (50 μM) treatment for 6 h. (D) Western blot analysis and quantification of phospho-JNK, JNK, phospho-ERK1/2, ERK1/2, phospho-p38, and p38 in primary LSECs pre-treated with NAC (5 mM) for 1 h and subsequently treated with H_2_O_2_ (100 μM) for 1 h. (E) *Bmp6* mRNA expression in primary LSECs pre-treated with U0126 (10 μM), SP600125 (5 μM), SB202190 (10 μM), or DMSO for 1 h, followed by H_2_O_2_ (100 μM) treatment for 6 h. All cell culture experiments in A, C, D, and E were conducted in the presence of hepatocyte-conditioned medium. Data in (B, C and E) are RT-qPCR results, normalized to *Rpl19* and presented as fold change compared to vehicle controls. The dashed line (ut) represents the mRNA expression of LSECs treated with the conditions shown, in the absence of H_2_O_2_. Data are obtained from three independent experiments and displayed as mean ± SD. The western blot in panels D shows a representative replicate of three independent experiments. Densitometric quantification displays mean ± SD of all biological replicates. Statistical significance: *p<0.05, **p<0.01, ***p<0.001, ****p<0.0001, one-way ANOVA, two-way ANOVA or Student’s t-test. CM, conditioned medium; FeNTA, ferric nitrilotriacetate; NAC, N-Acetyl-L-Cysteine.

Interestingly, H_2_O_2_ strongly activated the MAPK signaling pathway, as shown by increased phosphorylation of JNK, ERK1/2 and p38, an effect abolished by NAC (Figure 4D). To assess the role of MAPK signaling in H_2_O_2_-induced *Bmp6* mRNA expression, we used inhibitors targeting ERK1/2 (U0126), JNK (SP600125) and p38 (SB202190). We show that inhibition of the MAPK pathways abolishes H_2_O_2_-induced *Bmp6* expression (Figure 4E), suggesting that the oxidative stress-induced *Bmp6* response depends on MAPK activation. Consistently, H_2_O_2_-induced mRNA expression of *Hmox1* and *Nqo1* was also attenuated by MAPK inhibitors (Figure S3B).

Taken together, these findings demonstrate a critical role of the MAPK signaling pathway in controlling *Bmp6* expression in response to oxidative stress in LSECs.

## Discussion

Hepcidin is the master regulator of systemic iron fluxes. It inhibits dietary iron absorption and iron release from stores by binding to FPN expressed on duodenal enterocytes, macrophages and hepatocytes, respectively. Activation of hepcidin expression in hepatocytes by excess iron in the plasma and liver acts in a regulatory negative feedback loop to reduce iron absorption. Hepcidin transcription can be activated in hepatocytes by iron-bound transferrin involving iron sensing mechanisms mediated by the HC-associated proteins HFE, TFR2 and HJV, that directly or indirectly interact with the BMP receptors located on the surface of hepatocytes (1). In addition, hepcidin is an acute-phase protein whose transcription in hepatocytes is activated e.g. via IL-6 and the JAK-STAT signaling pathway (39), or by PAMPs via TLR2 and 6 (40). Interestingly, the hepatocyte-intrinsic mechanisms controlling hepcidin activation in response to iron, inflammation, hormonal, metabolic or oxidative stress signals are unable to function without active BMP/SMAD signaling (41-43), the major pathway controlling hepcidin gene expression (1). This requirement has been demonstrated in mice with deficiencies for the HC proteins HJV (44) and HFE (45), as well as for ALK3 (46) and SMAD4 (47), and in cell-based assays testing deletions of the BMP-response element in transfected hepcidin promoter constructs (41). The BMPs required for BMP/SMAD signaling in hepatocytes are produced by LSECs, with BMP6 playing a major role (8).

In this work we uncover that, in addition to iron-related signals, LSECs sense cell damage, redox and immune cues to activate *Bmp6* transcription via the MAPK signaling cascade. It is remarkable that similar signals not only control *Bmp6* in LSECs but also hepcidin in hepatocytes. This points towards an intensive cross-talk between the two cell types, which assures BMP/SMAD pathway activation in hepatocytes once hepcidin-activating external cues have been encountered. Signals controlling *Bmp6* expression in LSECs may thus determine the quantitative response of hepcidin by activating the BMP/SMAD signaling pathway in hepatocytes. Moreover, we hypothesize that the specificity of the BMP6-dependent hepcidin response to iron or inflammatory stimuli is dictated by hepatocytes (Figure 5).

**Figure 5.**
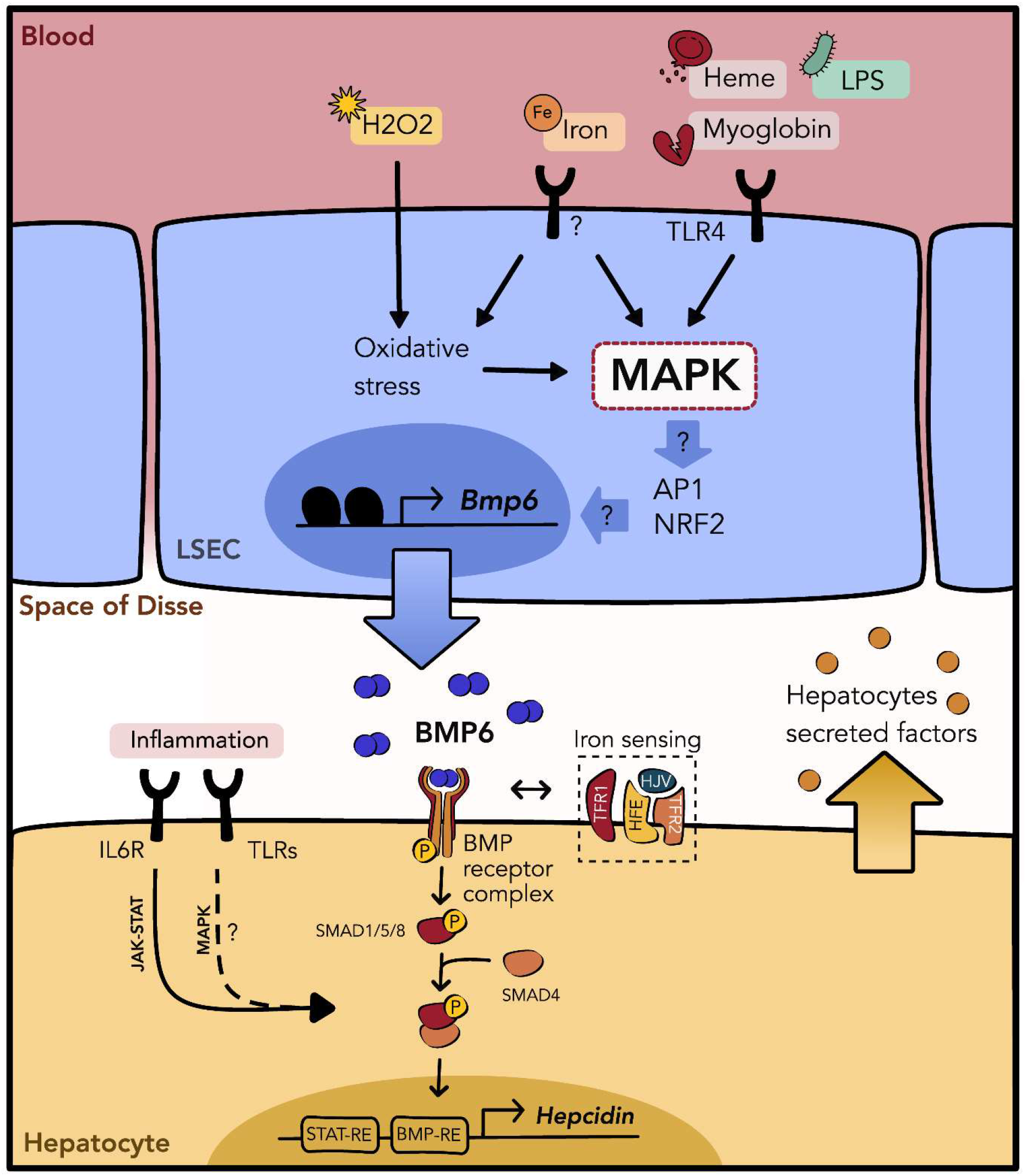
Working model of LSEC *Bmp6* regulation and hepcidin induction by iron, inflammatory and redox signals. In the presence of hepatocyte secreted factors, LSECs integrate iron, inflammatory (heme, myoglobin and LPS) and redox (H_2_O_2_) signals to induce *Bmp6* mRNA expression via the MAPK signalling pathway. While TLR4 mediates the response to inflammatory stimuli, the iron sensing receptor is still unknown. Most tested stimuli also activate the NRF2-mediated antioxidant response. However, how NRF2 interacts with the MAPK pathway for *Bmp6* regulation remains to be investigated. Upon its release in the space of Disse, BMP6 binds to the BMP receptor complex on the surface of hepatocytes, resulting in the activation of the SMAD signalling pathway. The active SMAD1/5/8-SMAD4 complex then binds to the BMP-responsive element (BMP-RE) in the promoter of the *Hepcidin* gene, allowing its transcription. Previous data have shown that *Hepcidin* induction by inflammatory stimuli via the JAK-STAT pathway requires a functional BMP-RE, in addition to the STAT-RE, in the *Hepcidin* promoter. Similarly, the hemochromatosis proteins HJV and HFE require active BMP signaling for iron-mediated *Hepcidin* induction in hepatocytes. Inflammatory cues can also activate hepcidin transcription via the TLRs-MAPK axis. However, a dependency on the BMP pathway has not been studied in this case. LPS, lipopolysaccharide; TLR, Toll-like receptor.

The integrative cross-talk between hepatocytes and LSECs is further supported by the observation that the *Bmp6* response to all tested stimuli is enhanced by the hepatocyte CM. The responsible factor(s) (22) has not yet been identified.

All immune and stress signals tested here that induce *Bmp6* mRNA expression do so by activating the MAPK signaling pathway in LSECs. While iron-related signals mainly activate the ERK1/2 branch, H_2_O_2_-induced oxidative stress and TLR4 stimulation by heme and LPS activate all three signaling branches, including ERK1/2, p38 and JNK. Formally, the MAPK pathway could be activated secondarily by non-canonical BMP signaling (48). However, we deem autocrine BMP signaling mechanisms as unlikely, as cycloheximide treatment did not alter *Bmp6* upregulation in response to LPS. In addition, our previous work demonstrated that in murine LSEC primary cultures autocrine responses to BMP2 and BMP6 are blocked despite high expression levels of the required receptor complexes (27).

The analysis of gene response patterns to heme treatment suggests that transcription factors downstream of the MAPK signaling (c-JUN, JUND, ELK1 and FOS) may act as mediators of heme-induced *Bmp6* expression. c-JUN, JUND and FOS are members of the AP-1 transcription factor family, which are well known MAPK pathway effectors working in (hetero)dimers. Their possible involvement in *Bmp6* transcription is in agreement with previous publications indicating c-JUN as iron-mediated *Bmp6* regulator (18).

Our findings also extend on previous studies that showed that iron can activate the p38/JNK branch of the MAPK pathway in LSECs, inducing *Bmp6* expression via the transcription factor ETS1 (19). Our data reveal that different branches of the MAPK pathway can mediate the *Bmp6* response. Differences may be explained by specifics of the model systems and treatment used as well as LSEC isolation and culturing methods. For example, Zurawska et al. showed that 1 mM ferric citrate activates p38 and *Bmp6* mRNA expression (19). By contrast, Fisher et al. reported that 200 μg/mL ferric ammonium citrate requires c-JUN to upregulate *Bmp6*, supporting a role of JNK. However, LSEC-specific *Jun*-KO mice do not show altered *Bmp6* and iron levels (18). Our results were obtained in primary murine LSEC cultures, which contain fenestrae (as observed *in vivo*), and express the LSEC-specific surface marker STAB2 (27).

It is also relevant to consider that the specificity of inhibitors of the various MAPK branches is limited (Figure 2E, F). Although different MAPK branches have been implicated as drivers of *Bmp6* induction, crosstalk frequently occurs among the canonical ERK, JNK, and p38 MAPK pathways (36). Consistently, the mTOR pathway, which was shown to regulate *Bmp6*, is also known to closely interact with the MAPK pathway (20, 49).

LSECs maintain hepatic immune responses and sustain immune tolerance by monitoring blood arriving from the portal vein that is continuously exposed to gut-derived antigens. LSECs express a wide array of TLRs that respond to a spectrum of ligands, triggering the production of pro-inflammatory cytokines, including TNF-α, IL-6, IL-1β, and IFN-β (13), which is also confirmed by our study (Figure S1H). We expect that pro-inflammatory cytokines, such as IL-6 and BMP6, are secreted under inflammatory conditions by LSECs. Such cytokines may act synergistically to enhance hepcidin expression in hepatocytes, explaining why the BMP/SMAD signaling pathway and BMP-RE in the hepcidin promoter are required for the inflammatory activation of hepcidin.

These findings are relevant in the context of hypoferremia and anemia of inflammation (AI). The persistent hepcidin induction observed in stress conditions, such as infection, iron overload, oxidative stress and hemolysis, contribute to hypoferremia development and AI, a widespread disorder caused by chronic inflammatory conditions (50). Here we show that PAMPs and DAMPs binding to TLR4 in LSECs upregulate *Bmp6*, an essential activator of hepcidin, suggesting that this mechanism may contribute to the generation of hypoferremia and AI.

The role of TLRs in activating *Bmp6* via MAPK signaling illuminates previous findings describing how TLRs expressed in macrophages recognize PAMPs and DAMPs and induce the production of inflammatory mediators (e.g. IL-6) that upregulate hepcidin in hepatocytes. More recently, TLR2/6 sensing in hepatocytes was further shown to directly induce *Hepcidin* mRNA expression via the MAPK signaling pathway (40). Taken together, these data suggest that PAMPs and DAMPs are sensed by several liver cell types via TLRs to control hepcidin and systemic iron homeostasis in a concerted manner. Future experiments are needed to validate our findings *in vivo*.

Our data fill an important knowledge gap by establishing LSECs as essential regulators of iron homeostasis during hemolysis, infection, and exposure to endogenous damage signals connecting innate immune sensing to the control of systemic iron homeostasis.

## Authors’ contribution

R.Q., M.W.H. and M.U.M. designed the experiments; R.Q., S.Cu., S.Co., C.M. performed the experiments; S.Co., S.A developed materials/techniques; R.Q., S.Cu., M.W.H. and M.U.M. drafted the manuscript, and all authors read and reviewed the manuscript; M.W.H. and M.U.M. supervised the project.

## Acknowledgment

M.U.M. acknowledges funding from the DFG (Priority Program SPP2306, GRK2727 and SFB1038), from the Federal Ministry of Education and Research (BMBF) (NephrESA project Nr 031L0191C), the Dietmar Hopp-Stiftung, and the Deutscher Akademischer Austauschdienst (A New Passage to India). M.U.M. and S.A. also acknowledge DFG-funding from FerrOs - FOR5146. S.Cu. is an associated PhD fellow of GRK2727.

