## Supplementary material for "Liver sinusoidal endothelial cells integrate metabolic and immune signals for MAPK-dependent BMP6 regulation and hepcidin induction"

### Supplementary methods

#### Isolation and culture of primary LSECs and hepatocytes

Primary mouse LSECs were isolated following a previously published protocol with minor modifications (1). Briefly, livers from C57BL/6N wild-type mice (22–25 weeks old) were perfused via the vena cava using Liver Perfusion Medium (Life Technologies #17701038), followed by enzymatic digestion with Liberase (Roche #5401127001). After mechanical decapsulation, the resulting liver cell suspension was collected in wash medium (Life Technologies #17704024) and sequentially filtered through 100 µm and 70 µm cell strainers. Hepatocytes and non-parenchymal cells (NPCs) were separated by low-speed centrifugation. Primary hepatocytes were seeded on collagen-coated plates at $2.5 \times 10^5$ cells/mL. NPCs were further enriched using density gradient centrifugation with Optiprep (Stemcell Technologies #07820). For primary LSEC culture, LSECs and KCs were separated by differential adhesion steps. LSECs were recovered and plated on collagen-coated plates at a density of $2 \times 10^6$ cells/mL. All primary cells were cultured in William's E medium (Life Technologies #32551020) supplemented with 4% fetal bovine serum (FBS) and 1% penicillin/streptomycin, under humidified conditions at 37 °C with 5% $CO_2$. Vascular endothelial growth factor (VEGF, Life Technologies #450-32) was additionally supplemented at the final concentration of 10 ng/mL. Hepatocyte-conditioned medium was prepared as previously described (2) and added to primary mouse LSECs 1 h before stimulation.

#### Cell culture treatments

Heme was used in the form of hemin and purchased from Frontiers Specialty Chemicals (#H651-9) and diluted in dimethyl sulfoxide (DMSO) according to the manufacturer's instructions to the indicated concentrations. The TLR4 inhibitor TAK242 (#13871), the MAPK pathway inhibitors RAF265 (#16991), U0126 (#70970), SB202190 (#21201) and Ulixertinib (#18298) were purchased from the Cayman Chemical Company and diluted in DMSO. Lipopolysaccharide (LPS) (#2630), myoglobin (#M1882), N-acetyl-L-cysteine (NAC) (#A7250), $H_2O_2$ (#1.07209) and cycloheximide (CHX) (#01810) were purchased from Sigma-Aldrich/Merck and diluted in phosphate buffered solution (PBS) (LPS, myoglobin, NAC and $H_2O_2$) or DMSO (CHX). The TLR ligands Pam3CSK4 (TLR2:1) (#tlrl-pms), PGN - SA (TLR2) (#tlrl-pgns2), Poly I:C (TLR3) (#tlrl-picw), FLA-ST (TLR5) (#tlrl-stfla), FSL1 (TLR2:TLR6) (tlrl-fsl), R848 (TLR7:8) (#tlrl-r848-1), ODN (TLR9) (#tlrl-1826) and the MAPK inhibitor SP600125 (#tlrl-sp60) were purchased form Invivogen and diluted in PBS (TLR ligands) or DMSO (SP600125). Ferric nitrilotriacetate (FeNTA) was prepared dissolving iron (III) chloride hexahydrate (Sigma-Aldrich/Merck #31232) and nitrilotriacetic acid disodium salt (NTA) (Sigma-Aldrich/Merck #N0128) in ratio 1:4 in PBS.

**Protein extraction**

Total protein lysates were prepared by homogenizing cell pellets in radioimmunoprecipitation assay (RIPA) buffer supplemented with a protease inhibitor cocktail (cOmplete™, Roche #04693116001) and phosphatase inhibitors (PhosSTOP™, Roche #04906845001). Protein concentrations were determined using the bicinchoninic acid (BCA) assay (Life Technologies #23228, #23224). Equal amounts of protein (20 µg per sample) were loaded for SDS-PAGE and Western Blot.

**Reactive oxygen species (ROS) measurement**

Intracellular ROS levels in LSECs were detected using the CellROX® Green Flow Cytometry Assay Kit (Life Technologies #C10492) according to the manufacturer's instructions. Following treatments, cells were incubated with 500 nM CellROX® Green for 30 minutes, then stained with 5 nM SYTOX® Red Dead Cell Stain (Life Technologies #S34859) for 5 minutes. Fluorescence intensity and the percentage of ROS-positive, live cells were measured using a BD Accuri™ C6 Plus flow cytometer.

**RNA sequencing (RNA-seq) data analysis**

Reads were aligned to the GRCm39 genome using STAR aligner (2.7.11b) (3). Transcription factor inference analysis was conducted using the CollecTRI (Collection of Transcriptional Regulatory Interactions) resource with the univariate linear model available from the decoupleR (v2.14.0) package (4). Gene Set Enrichment Analysis (GSEA) was performed using the GSEA software 4.4.0 (5). Differential expression analysis has been performed using the R library DESeq2 (v1.48.1) (6) and statistics (*) refers to p-adjusted values.

Supplementary tables

**Table S1: List of antibodies and dilutions**

| Target protein | Dilution ratio | Source | Provider | Catalog No. |
|---|---|---|---|---|
| Mouse IgG | 1:5000 | Rabbit | Sigma-Aldrich/Merck | A9044 |
| Rabbit IgG | 1:5000 | Goat | Sigma-Aldrich/Merck | A0545 |
| AKT | 1:1000 | Rabbit | Cell Signaling | 9272S |
| ERK1/2 | 1:1000 | Rabbit | Cell Signaling | 4695 |
| JNK | 1:1000 | Rabbit | Cell Signaling | 9252 |
| p38 MAPK | 1:1000 | Rabbit | Cell Signaling | 8690 |
| Phospho-AKT (Ser473) | 1:1000 | Rabbit | Cell Signaling | 4060 |
| Phospho-ERK1/2 (Thr202/Tyr204) | 1:1000 | Rabbit | Cell Signaling | 4370 |
| Phospho-JNK (Thr183/Tyr185) | 1:1000 | Rabbit | Cell Signaling | 4668 |
| Phospho-p38 MAPK (Thr180/Tyr182) | 1:1000 | Rabbit | Cell Signaling | 4511 |
| Vinculin | 1:1000 | Mouse | Sigma-Aldrich/Merck | SAB4503069 |

**Table S2 List of RT-qPCR primers**

| **Target gene** | **Forward primer 5’- 3’** | **Reverse primer 5’- 3’** |
|---|---|---|
| *Bmp6* | ATGGCAGGACTGGATCATTGC | CCATCACAGTAGTTGGCAGCG |
| *Ccnd1* | GCAGAAGGAGATTGTGCCATCC | AGGAAGCGGTCCAGGTAGTTCA |
| *Hmox1* | AGGCTAAGACCGCCTTCCT | TGTGTTCCTCTGTCAGCATCA |
| *Il6* | GCTACCAAACTGGATATAATCAGGA | CCAGGTAGCTATGGTACTCCAGAA |
| *Nos2* | GAGACAGGGAAGTCTGAAGCAC | CCAGCAGTAGTTGCTCCTCTTC |
| *Nqo1* | AGCGTTCGGTATTACGATCC | AGTACAATCAGGGCTCTTCTCG |
| *Rpl19* | AGGCATATGGGCATAGGGAAGAG | TTGACCTTCAGGTACAGGCTGTG |
| *Tnfα* | *TGCCTATGTCTCAGCCTCTTC* | *GAGGCCATTTGGGAACTTCT* |

Supplementary results

### Figure legends:

#### Figure S1. TLR4 activation by LPS, heme, and myoglobin drives pro-inflammatory cytokine production

(A) Expression profile of *Tlr1–9* mRNA in primary cultured LSECs (n = 3). (B) *Bmp6* mRNA expression in primary LSECs treated with myoglobin (50 µM) or vehicle control (ut, untreated) for 6 h. (C-E) *Il6* mRNA expression in LSECs pre-treated with TAK242 (5 µM) for 1 h, followed by LPS (5 ng/mL), heme (2.5 µM), or myoglobin (50 µM) treatment for 6 h. (F) *Bmp6* mRNA expression in LSECs pre-treated with TAK242 (5 µM) for 1 h, followed by myoglobin (50 µM) treatment for 6 h. (G) Gene set enrichment analysis (GSEA) was performed using the WikiPathways database to identify biological pathways enriched in heme-treated compared to untreated control LSECs. The dotplot shows significant (FDR < 25%) enriched pathways upon LSECs treatment. The normalized enrichment score (NES) indicates the degree to which a pathway is overrepresented in the heme-treated group. The size of the dots represents the number of genes contributing to each pathway, while the color gradient corresponds to the nominal (NOM) p-value. (H) Heatmap showing significant (adjusted p-value < 0.1) differentially expressed cytokine gene levels in RNA-seq from LSECs treated with heme vs untreated controls. Values are represented as Z-score of log2-transformed expression levels (variance-stabilized transformation from DeSeq2 package) across samples. (I) *Il6* and *Tnfa* mRNA expression in LSECs treated with 5 ng/mL LPS for 6 h. (J) *Nos2* mRNA expression in primary LSECs pre-treated with CHX (5 µM) or DMSO for 1 h, followed by 5 ng/mL LPS treatment for 6 h. (K) Heatmap showing significant (adjusted p-value < 0.1) differentially expressed LSEC marker gene levels in RNA-seq from LSECs treated with heme vs untreated controls. Values are represented as Z-score of log2-transformed expression levels (variance-stabilized transformation from DeSeq2 package) across samples. (L) *Hmox1* and *Nqo1* mRNA expression in LSECs treated with 2.5 µM heme for 6 h. (M) *Hmox1* mRNA expression in LSECs treated with myoglobin (50 µM) for 6 h. (N) *Hmox1* and *Nqo1* mRNA expression in LSECs treated with 5 ng/mL LPS. (O) *Nqo1* mRNA expression in primary LSECs pre-treated with U0126 (10 µM), SP600125 (5 µM), SB202190 (10 µM), or DMSO for 1 h, followed by 2.5 µM heme treatment for 6 h. All cell culture experiments were conducted in the presence of hepatocyte-conditioned medium. Gene expression levels were assessed by RT-qPCR, normalized to the housekeeping gene *Rpl19*, and expressed as fold change relative to vehicle-treated controls. The dashed line (ut) represents the mRNA expression of LSECs treated with the conditions shown, in the absence of LPS, heme or myoglobin. Data are obtained from three independent experiments and displayed as mean ± SD. Statistical significance: *p<0.05, **p<0.01, ***p<0.001, one-way ANOVA or Student's t-test. CHX, cycloheximide; TPM, transcripts per million; FDR, false discovery rate.

**Figure S2. Inhibition of the MAPK signaling pathway decreases FeNTA-induced NRF2 target gene expression.**

(A) *Ccnd1* mRNA expression in primary LSECs pre-treated with RAF265 (5 μM), U0126 (10 μM), ulixertinib (10 μM), or DMSO for 6 h. (B) *Hmox1* and *Nqo1* mRNA expression in primary LSECs pre-treated with RAF265 (5 μM), U0126 (10 μM), ulixertinib (10 μM), or DMSO for 1 h, followed by 50 μM FeNTA treatment for 6 h. All experiments were performed in the presence of hepatocyte-conditioned medium. Data are RT-qPCR results, normalized to the housekeeping gene *Rpl19* and presented as fold change relative to vehicle controls. The dashed line (ut) indicates the mRNA expression of LSECs treated with DMSO or the inhibitors in the absence of FeNTA. Data are derived from three independent experiments and presented as mean ± SD. Statistical significance: *p<0.05, **p<0.01, ***p<0.001, ****p<0.0001, one-way ANOVA. FeNTA, ferric nitrilotriacetate.

**Figure S3. MAPK signaling is involved in oxidative stress-mediated NRF2 activation.**

(A) *Hmox1* and *Nqo1* mRNA expression in primary LSECs pre-treated with NAC (5 mM) for 1 h, followed by treatment with hydrogen peroxide ($H_2O_2$, 100 μM), FeNTA (50 μM) or vehicle control (NT, non-treated) for 6 h. (B) *Hmox1* and *Nqo1* mRNA expression in primary LSECs pre-treated with U0126 (10 μM), SP600125 (5 μM), SB202190 (10 μM), or DMSO for 1 h, followed by 50 μM $H_2O_2$ treatment for 6 h. All experiments were performed in the presence of hepatocyte-conditioned medium. Data are RT-qPCR results, normalized to the housekeeping gene *Rpl19* and presented as fold change relative to vehicle controls. The dashed line (ut) indicates the mRNA expression of LSECs treated with DMSO or the inhibitors in the absence of $H_2O_2$. Data are derived from three independent experiments and presented as mean ± SD. Statistical significance: *p<0.05, **p<0.01, ***p<0.001, ****p<0.0001, one-way ANOVA or two-way ANOVA. FeNTA, ferric nitrilotriacetate; NAC, N-acetyl-L-cysteine.

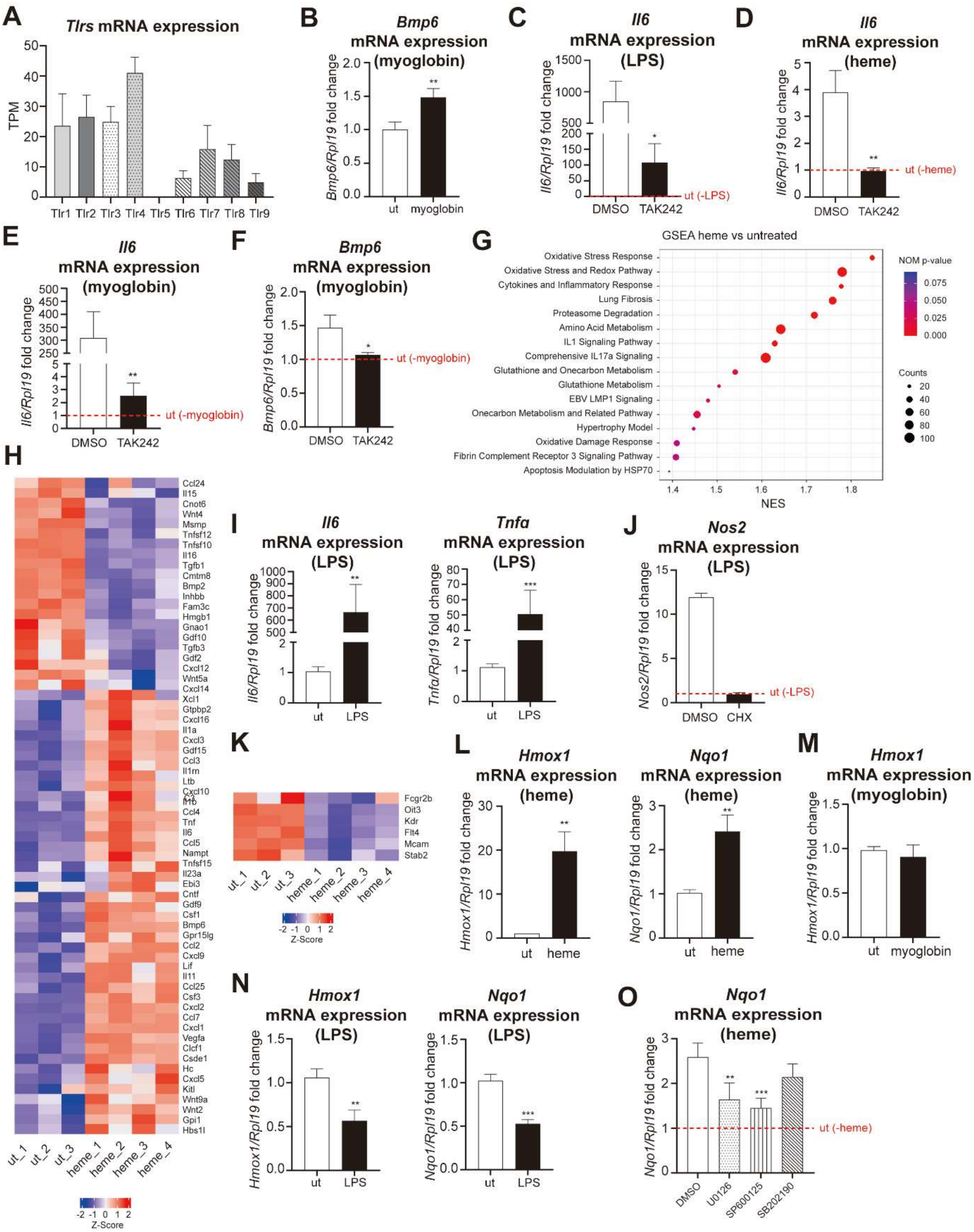


Supplementary figure 1

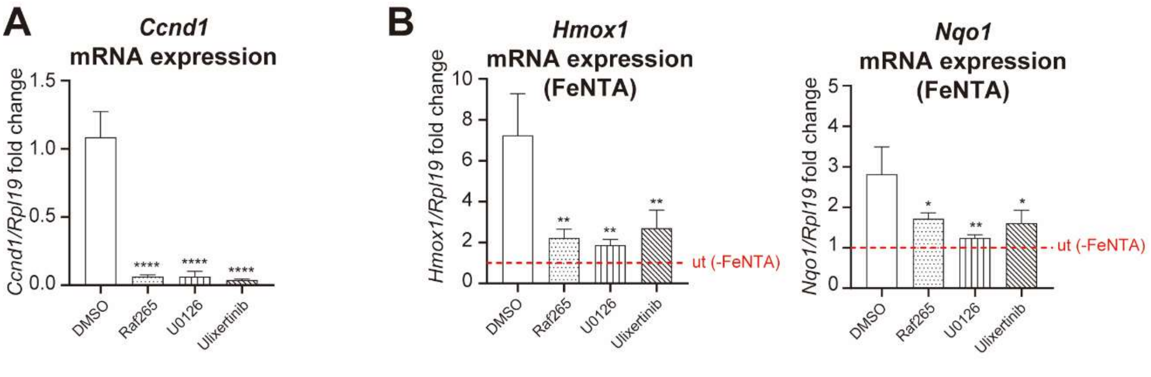


Supplementary figure 2

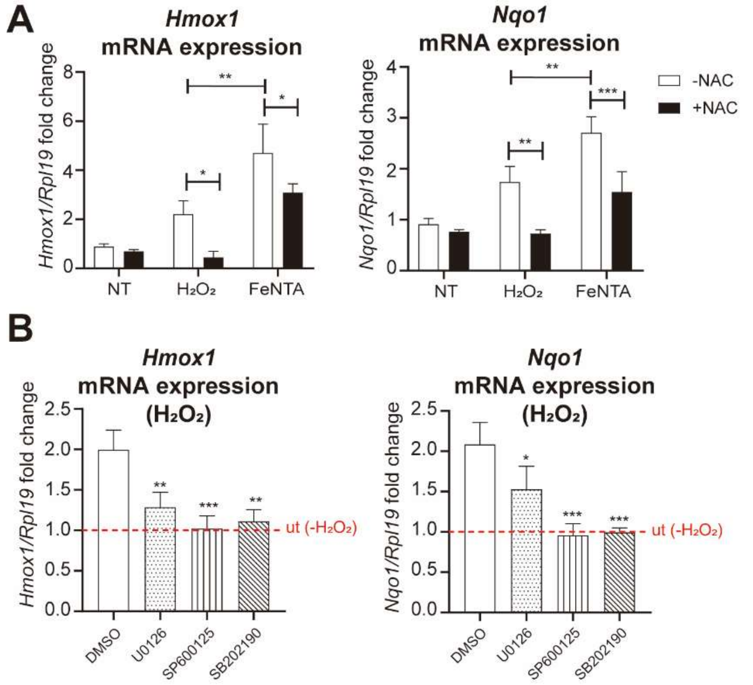


Supplementary figure 3
